# Paraquat Licences α-Synuclein to Trigger NLRP3-Dependent Pyroptosis in Microglia

**DOI:** 10.1101/2025.07.23.666340

**Authors:** Yuzhihan Cao, Vandana Deora, Nemat Khan, Avril A. B. Robertson, Kate Schroder, Matthew A. Cooper, Trent M. Woodruff, Eduardo A. Albornoz

## Abstract

Parkinson’s disease (PD) is characterized by dopaminergic neurodegeneration and the pathological accumulation of α-synuclein (αSyn) aggregates. While its etiology is multifactorial, environmental toxicants such as paraquat, a widely used herbicide banned in over 70 countries but still permitted in some regions, including Australia, are strongly implicated. We previously showed that αSyn activates the NLRP3 inflammasome in microglia, inducing interleukin-1β (IL-1β) release without triggering pyroptosis. Here, we examine how paraquat modulates microglial NLRP3 activation by αSyn. Using primary microglia from wild-type (WT) and *Nlrp3* knockout (*Nlrp3*^-/-^) mice, alongside human monocyte-derived microglia, we demonstrate that paraquat induces dose-dependent NLRP3 activation in LPS-primed WT microglia via NADPH oxidase-mediated reactive oxygen species (ROS) generation. This results in ASC speck formation and IL-1β release, which is absent in *Nlrp3*^-/-^ microglia and blocked by the selective NLRP3 inhibitor MCC950. While αSyn alone fails to induce pyroptosis, co-treatment with paraquat resulted in caspase-1-dependent lactate dehydrogenase (LDH) release and gasdermin-D cleavage, hallmarks of pyroptotic cell death. This response was also suppressed by NLRP3 inhibition. Our findings reveal that paraquat acts as a critical second hit to license αSyn-induced inflammasome-dependent pyroptosis in microglia, highlighting a mechanistic link between environmental toxin exposure and innate immune activation in PD pathogenesis. Targeting this axis with brain-penetrant NLRP3 inhibitors may offer an alternative therapeutic avenue for synucleinopathies associated with environmental toxins.

## Introduction

Parkinson’s disease (PD) is a progressive neurodegenerative disorder characterized by chronic neuroinflammation, which plays a central role in dopaminergic neuronal loss and the overall pathophysiology of the disease (Tansey et al., 2022). Although the etiology of PD remains largely unknown, several disease-associated genes and environmental risk factors have been identified as modulators of immune function. A key pathogenic feature is the accumulation of α-synuclein (αSyn) aggregates, a major component of Lewy bodies and a hallmark neuropathological feature of PD (Luk et al., 2012; Obeso et al., 2010).

Among environmental risk factors, chronic exposure to pesticides and herbicides has been consistently linked to increased PD risk. Paraquat (N, N′-dimethyl-4,4′-bipyridinium dichloride), a widely used broad-spectrum herbicide, has been associated with PD in multiple epidemiological studies (Costello, Cockburn, Bronstein, Zhang, & Ritz, 2009; Lee, Bordelon, Bronstein, & Ritz, 2012; Liou et al., 1997; Sharma & Mittal, 2024; Tanner et al., 2011).

In Australia, the continued use of paraquat has raised significant concern due to its neurotoxic effects, particularly as it remains approved for certain agricultural applications. This persistence is largely driven by economic considerations, with a potential ban predicted to increase cropping costs (Alison Walsh, 2021), despite paraquat being prohibited in many other countries. Chronic exposure to paraquat has been shown to reproduce key pathological features of PD, including dopaminergic neurodegeneration (Betarbet et al., 2000). Experimental evidence further supports these concerns, indicating that paraquat can directly damage dopaminergic neurons (Richardson, Quan, Sherer, Greenamyre, & Miller, 2005) or exert its neurotoxicity primarily through the generation of reactive oxygen species (ROS) and microglial activation (Wu et al., 2005).

These risk factors converge on a central hallmark of neurodegenerative diseases: neuroinflammation. Increasing evidence implicates activation of the brain’s innate immune system in this process (Heneka, Kummer, & Latz, 2014), specifically via the activation of the NOD-like receptor family pyrin domain-containing 3 (NLRP3) inflammasome in microglia (Song, Pei, Yao, Wu, & Shang, 2017).

The NLRP3 inflammasome is an intracellular multiprotein complex responsible for the maturation of proinflammatory cytokines such as interleukin-1β (IL-1β), and for triggering a highly inflammatory form of cell death termed pyroptosis, mediated by caspase-1 and the adaptor protein ASC (apoptosis-associated speck-like protein containing a CARD) (Gross, Thomas, Guarda, & Tschopp, 2011). Our previous work demonstrated that the NLRP3 inflammasome is upregulated in the brains of post-mortem PD patients and in preclinical mouse models, specifically within microglia. We further confirmed these findings *in vitro,* showing that pre-formed α-synuclein fibrils (αSyn) activate NLRP3 in microglial cells, resulting in IL-1β release without concomitant pyroptosis (Albornoz, Gordon, et al., 2023; Gordon et al., 2018).

In parallel, paraquat has been shown to induce IL-1β secretion via the NLRP3-ASC-caspase-1 pathway in peripheral macrophages, a mechanism implicated in paraquat-induced acute lung injury (Z. Liu et al., 2015). However, it remains unclear whether paraquat activates the NLRP3 inflammasome in microglia, either alone or in combination with αSyn, under conditions of exposure in the context of synucleinopathy, and how this contributes to PD pathogenesis. In this study, we investigated the interplay between paraquat, αSyn, and microglia in the context of NLRP3 inflammasome activation. Using primary microglial cultures derived from wild-type (WT) and *Nlrp3* knockout mice (*Nlrp3*^-/-^), as well as human monocyte-derived microglial-like cells (MDMi) (Albornoz, Amarilla, et al., 2023), we demonstrate that paraquat triggers NLRP3 inflammasome activation in LPS-primed microglia via an NADPH-oxidase-dependent ROS pathway. This activation culminates in pyroptotic cell death, exacerbated by the presence αSyn, underscoring a synergistic effect that may exacerbate neuroinflammation and neurodegeneration in PD.

## Materials and methods

### Chemicals and reagents

Paraquat (1, 1’-dimethyl-4, 4’-bipyridinium dichloride), anti-β-tubulin antibody (T4026) and the non-selective NADPH inhibitor diphenylene iodonium (DPI) were purchased from Sigma-Aldrich Inc., St. Louis, MO. The NADPH oxidase NOX2 inhibitor GSK2795039 was obtained from Selleck Chemicals USA. DMEM/F-12 Cat #11320, heat-inactivated fetal bovine serum, L-glutamine, IR-dye tagged secondary antibodies, penicillin, and streptomycin and other cell culture reagents was obtained from Invitrogen (Gaithersburg, MD). The specific caspase-1 inhibitor VX-765 and Ultrapure LPS (E.Coli 0111: B4) was obtained from Invivogen. Antibodies for mouse NLRP3 (Cryo-2), ASC (AL177) and caspase-1 (Casper-1) were obtained from Adipogen International. Goat polyclonal antibody to IL-1β was purchased from R&D systems.

### Preparation of fibrillar synuclein aggregates

Recombinant human synuclein was obtained from rPeptide Inc. (Bogart, GA) and *in vitro* fibril generation was performed by incubation at 37°C with agitation in an orbital mixer (400rpm) during a week with daily sonication until the day of treatment (Albornoz, Gordon, et al., 2023). The generation of fibrillar Syn species was confirmed by transmission electron microscopy and Thioflavin T fluorescence before use.

### Primary mouse microglia cultures

The animal ethics committee approved all animal procedures at The University of Queensland. Primary microglial cultures were prepared from C57BL/6 or *Nlrp3*^-/-^ postnatal day 1 (P1) mouse pups and purified by column free magnetic separation system as described previously (Deora et al., 2020). Primary microglia cultures were maintained in DMEM/F12 complete medium (DMEM-F12, GIBCO supplemented with 10% heat-inactivated FBS, 50 U/mL penicillin, 50 μg/mL streptomycin, 2 mM L-glutamine, 100 μM non-essential amino acids, and 2 mM sodium pyruvate). And the cells were then incubated in 5% CO2 incubator at 37 °C until further treatments.

### Primary microglia treatment paradigm for inflammasome activation

For inflammasome activation experiments, 1 x 10^5^ cells of wild type (WT) or *Nlrp3*^-/-^ primary microglia were plated in 96 well plate and 5 x 10^5^ in 12 well plate with serum-free media overnight and primed with 200 ng/ml of ultrapure LPS for 3 h. Cells were then washed in a serum-free medium after priming to remove any residual LPS and stimulated with conventional NLRP3 inflammasome activators ATP (5 mM) or Syn fibrils (5 μM) for the indicated time points. Dose-response studies cells were stimulated with paraquat (50, 100, 200, 400 μM) at 24 h. Time course studies cells were stimulated with paraquat 100 μM for 4, 18, 24 h. For inhibition studies, all compounds including VX-765 (20 μM), GSK2795039 (20, 40 μM) diphenylene iodonium (DPI, 5-50 nM) and MCC950 (100 nM) were added at the end of the priming step 30 minutes followed by activation. At the end of treatment, the supernatants were collected and stored at -80°C until analysis by ELISA, LDH assay or western blotting.

### Generation of human Monocyte-Derived Microglia (MDMi)

Ethical approval for collecting and utilising human donor blood was obtained from The University of Queensland Human Research Ethics Committee. Monocytes were isolated from buffy coats obtained from Australian Red Cross Lifeblood as we previously described (Albornoz, Amarilla, et al., 2023). Briefly, buffy coat was diluted 1:1 with phosphate buffered saline (PBS) 2% fetal bovine serum (FBS) and transferred into sterile SepMate 50 (STEMCELL Technologies, BC, Canada) as per manufacturer’s instructions. Peripheral blood monocytes (PBMCs) were collected. Monocytes were positively selected from whole PBMCs using anti CD14+ microbeads (Miltenyi Biotec) and plated at the following densities per well: 1 × 10^5^ cells (96-well plate) and 3 × 10^5^ cells (24-well plate). To induce the differentiation of MDMi, we incubated monocytes under serum-free conditions using RPMI-1640 Glutamax (Life Technologies) with 1% penicillin/ streptomycin (Lonza) and Fungizone (2.5 μg/ml; Life Technologies) and a mixture of the following human recombinant cytokines: M-CSF (10 ng/ml; Preprotech, 300-25), GM-CSF (10 ng/ml; Preprotech, 300-03), NGF-β (10 ng/ml; Preprotech, 450-01), MCP-1(CCL2) (100 ng/ml; Preprotech, 300-04), and IL-34 (100 ng/ml; Preprotech, 200-34-250) under standard humidified culture conditions (37 °C, 5% CO2) for up to 14 days. Differentiation of PBMCs into MDMi was confirmed by immunofluorescence for microglial markers.

### Western blotting and ELISA

At the end of each treatment supernatants were collected and proteins were concentrated by the addition of an equal volume of methanol-chloroform precipitation as previously described (Jakobs, Bartok, Kubarenko, Bauernfeind, & Hornung, 2013). The microglial cell lysates were prepared using RIPA buffer (Pierce). The protein samples were separated using Bio-Rad 4-20% precast gels and then transferred onto nitrocellulose membranes. Membranes were blocked in the Licor blocking buffer for 1 hr. Primary antibodies for NLRP3, and Caspase-1 were prepared by at a concentration of 1:1,000 as specified by the manufacturer. After incubation with Licor IR dye-labelled secondary antibodies (1:10,000) and washing steps, the membranes were scanned using Licor Odyssey CLX imaging system. For densitometry analysis, band intensities were normalized against the GAPDH loading control and represented as the fold change over vehicle-treated controls. For ELISA analysis, the mouse IL-1β kit R&D Systems Cat # DY008, and Human IL-1β kit R&D Systems (Cat# DY201) was used to measure IL-1β levels in the supernatant of activated microglia following the manufacturer instructions.

### Immunocytochemistry

Primary WT and *Nlrp3*^-/-^ microglia and MDMi were grown on poly-D lysine coated coverslips and at the end of treatment were fixed for 15 minutes by adding an equal volume of 4% PFA into the treatment media. Cells were then washed with PBS several times and incubated in blocking buffer (PBS containing 2% BSA, 0.1% Triton-X and 0.05% Tween) for 1 h at room temperature. Coverslips were then incubated overnight at 4°C overnight with primary antibodies for ASC (1: 500, Adipogen) and β-tubulin (1:2,000, Sigma). Antibodies were diluted in PBS containing 2% BSA. Samples were then washed several times with PBS and incubated with Alexa dye-conjugated secondary antibodies. The nucleus was labeled with DAPI stain and coverslips were mounted with Prolong Gold antifade mounting medium.

### Quantification of Caspase-1 mediated pyroptosis

At the end of each treatment, supernatants were collected and LDH release was quantified using an LDH assay kit (TOX7, Sigma) as per the manufacturer’s instructions. Caspase-1 dependent LDH release that was inhibited by the highly-specific small molecule caspase-1 inhibitor VX-765 (20 μM) was used as a read-out for pyroptosis, and compared to nigericin (10 μM) as positive control, we previously described (Deora, Albornoz, Zhu, Woodruff, & Gordon, 2017)

### Membrane permeability assay

SYTOX Green (Life Technologies) (0.5 mM in Serum free media) was added to primary microglia (1 x10^6^ /well) at the end of each treatment for 10 minutes. Dye uptake was measured on a microplate reader (485/530 nm) to assess membrane permeability.

### Data Analysis

Statistical analysis was performed using GraphPad Prism 10.4 software. Data are represented as mean ± S.E.M. from at least 3 independent experiments. ANOVA followed by Tukey’s post-test was performed to compare all treatment groups. ns (non-significant), *** P < 0.001 and **** P < 0.0001 denote statistically significant differences between indicated group

## Results

### Paraquat activates the NLRP3 inflammasome in primary mouse microglia via a NOX2-dependent ROS pathway

To determine whether paraquat activates the NLRP3 inflammasome in microglia, we measured IL-1β release in the supernatant of primary wild-type (WT) and *Nlrp3* knockout (*Nlrp3*^-/-^) mouse microglia (Fig. 1B-E). Cells were first primed with lipopolysaccharide (Vehicle; LPS 200 ng/ml) for 3h (Fig1A), followed by stimulation with increasing concentrations of paraquat. LPS-primed WT microglia showed a significant dose- (50, 100, 200, and 400 μM) and time-dependent (4, 18, and 24 h; 100 μM) increase in IL-1β release. ATP (5 mM) served as a positive control for NLRP3 inflammasome activation (Fig. 1B,D), as previously described (Zha et al., 2016). Notably, IL-1β release was absent in *Nlrp3*^-/-^ microglia treated with either paraquat or ATP, confirming that this response is NLRP3-dependent (Fig. 1C, E).

**Figure 1.**
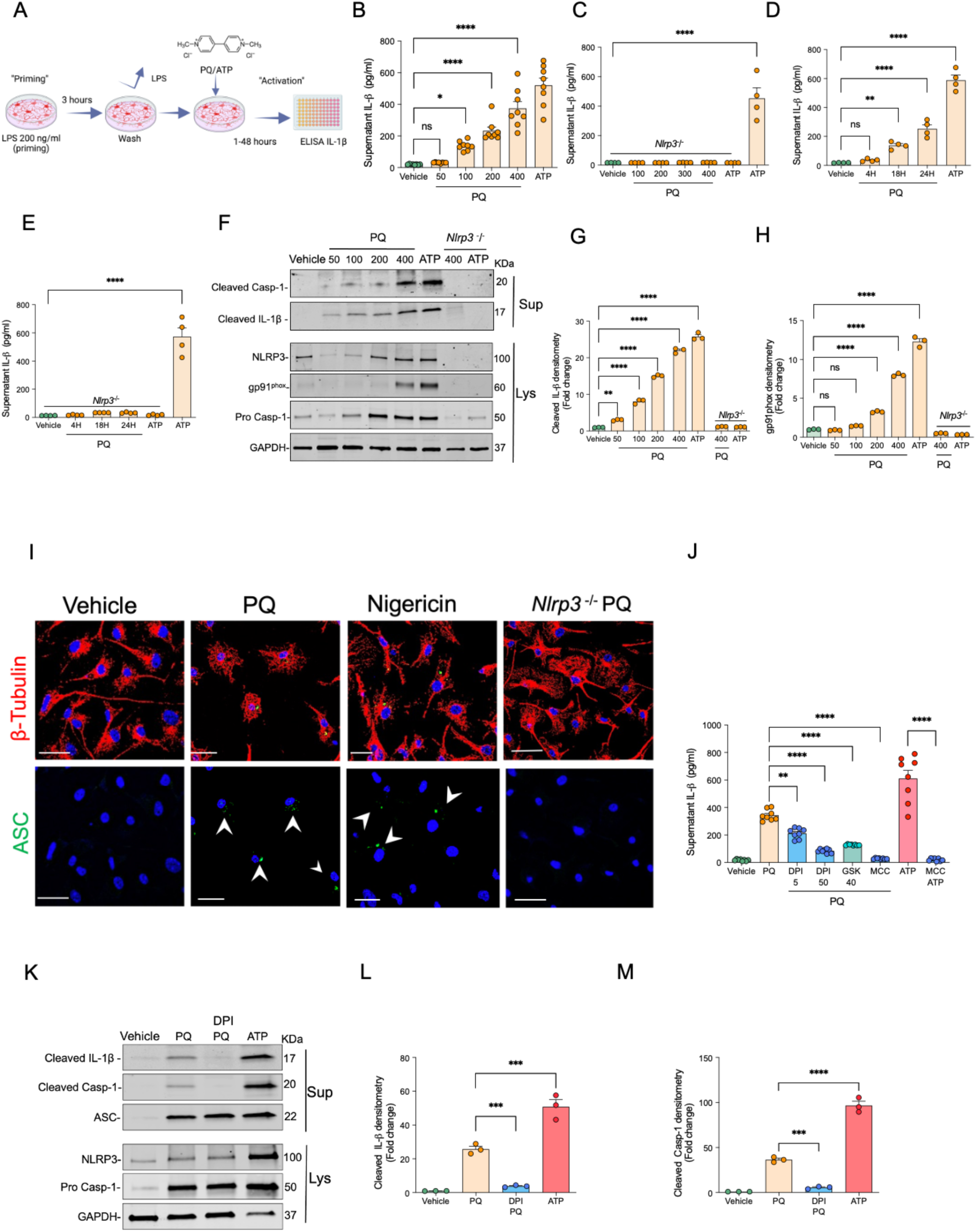
Paraquat activates the NLRP3 inflammasome in primary microglia via NOX2- dependent ROS pathway. **(A)** Schematic of the experimental paradigm: LPS-primed primary microglia were exposed to paraquat (PQ) **(B)** Dose-dependent IL-1β secretion in LPS-primed (vehicle) primary microglia treated with increasing concentrations of PQ (50,100,200,400 μM) for 24h ATP (5 mM, 1 h) served as a positive control **(C)** IL-1β secretion in Nlrp*3* ^-/-^ microglia treated under the same conditions. **(D-E)** Time course of IL-1β secretion following exposure to PQ (200 μM) for 24h in WT (D) and Nlrp*3* ^-/-^ **(E)** microglia. **(F)** Immunoblot analysis of cleaved caspase-1 (p20) and cleaved IL-1β (p17) in supernatants (Sup), and NLRP3, and pro caspase-1 in cell lysates (Lys) from WT and *Nlrp3* ^-/-^ microglia treated with PQ (50–400 μM) for 24 h. **(G-H)** Densitometric quantification of cleaved IL-1β (p17) and gp91^phox^ levels from western blots (I) Immunofluorescence staining for ASC (green) in WT and *Nlrp3* ^-/-^ microglia following PQ (200 μM, 24h), showing ASC speck formation (white arrows). Nigericin (10 μM) was used as a positive control. Nuclei were counterstained with DAPI (blue). Scale bar: 30 μm. (J) IL-1β secretion in WT microglia exposed to PQ (200 μM, 24h), in the presence of diphenyleneiodonium (DPI, 5 or 50 nM), the NOX2-specific inhibitor GSK2795039 (40 μM), or the NLRP3 inhibitor MCC950 (100 nM) (K) Immunoblot analysis of cleaved IL-1β (p17), cleaved caspase-1 (p20), and ASC in supernatants, and NLRP3 and pro–caspase-1 in lysates from WT microglia pretreated with DPI (50 nM) (L-M) Densitometric quantification of cleaved IL-1β (p17) and) and cleaved caspase-1 (p20) levels from western blots Data are shown as mean ± SEM from ≥3 independent experiments. Statistical significance by one-way ANOVA with Tukey’s post hoc test; \**P* < 0.05 ****p < 0.0001; ns, not significant.

To further assess inflammasome activation, we performed western blot analysis of cell supernatants (Sup) and lysates (Lys) (Fig.1F-H). Paraquat treatment led to a dose-dependent increase in cleaved caspase-1 (p20) and cleaved IL-1β (p17) in the supernatants, along with upregulated NLRP3, pro-caspase 1(p50) and NADPH oxidase 2 (NOX2, gp91^phox^) in WT microglia lysates. As expected, these components were absent or markedly reduced in *Nlrp3*^-/-^ microglia.

We visualized inflammasome assembly by immunostaining for ASC specks (Albornoz, Amarilla, et al., 2023; Yu et al., 2023). Paraquat-treated WT microglia exhibited characteristic ASC speck formation (green; Fig. 1I), similar to nigericin-treated positive controls (Gluck et al., 2023). In contrast, ASC specks were not observed in *Nlrp3*^−/−^ microglia following paraquat treatment, further confirming NLRP3 dependence.

Paraquat triggers reactive oxygen species (ROS) production via a well-characterized redox cycling mechanism in which NADPH oxidase plays a central role (Castello, Drechsel, & Patel, 2007; Cristovao, Choi, Baltazar, Beal, & Kim, 2009). ROS is a known activator of the NLRP3 inflammasome (Abais, Xia, Zhang, Boini, & Li, 2015; Dominic, Le, & Takahashi, 2022). To investigate whether this mechanism underlies paraquat-mediated inflammasome activation, we used diphenyleneiodonium (DPI), a non-selective NADPH oxidase inhibitor previously shown to block paraquat-induced ROS (Miller, Sun, & Sun, 2007). DPI (5 and 50 nM) significantly reduced IL-1β release in paraquat-treated microglia when added after LPS priming. Similarly, the NOX2-specific inhibitor GSK2795039 (Hirano et al., 2015) (40 μM) attenuated IL-1β release, implicating NOX2-derived ROS in paraquat-mediated inflammasome activation (Surace & Block, 2012; Taetzsch & Block, 2013). As a control for NLRP3 inhibition, MCC950 completely abolished IL-1β release in response to paraquat and ATP (Fig. 1J).

Western blot analysis further revealed that DPI suppressed the release of cleaved IL-1β (p17) and caspase-1 (p20) suggesting that ROS is essential for inflammasome activation.

### Paraquat enhances fibrillar α-synuclein–induced NLRP3 activation via NOX2-derived ROS

Lewy body inclusions, rich in misfolded α-synuclein (αSyn), are a hallmark of PD (Peelaerts et al., 2015; Uemura et al., 2023). We previously demonstrated that fibrillar α-synuclein (αSyn,10 μM) induces NLRP3 inflammasome activation in LPS-primed microglia after 24h (Albornoz, Gordon, et al., 2023; Gordon et al., 2018). Having established paraquat as an NLRP3 activator, we next examined whether paraquat enhances αSyn-induced inflammasome activation.

Microglia were exposed to paraquat (PQ; 100 μM), αSyn (5 μM), or both (PQ+Syn), and IL-1β levels were measured in the supernatant (Fig. 2A). Both paraquat and αSyn alone significantly increased IL-1β release compared to LPS-primed vehicle controls with paraquat+αSyn co-treatment producing a ∼3-fold increase relative to either treatment alone (Fig. 2B). This enhancement was abolished by pre-treatment with the selective NLRP3 inhibitor MCC950 (100 nM), confirming NLRP3 involvement. Western blot analysis showed increased cleaved IL-1β (p17), cleaved caspase-1 (p20), and NLRP3 levels in paraquat+αSyn- treated cells compared to single treatments (Fig. 2C-E).

**Figure 2.**
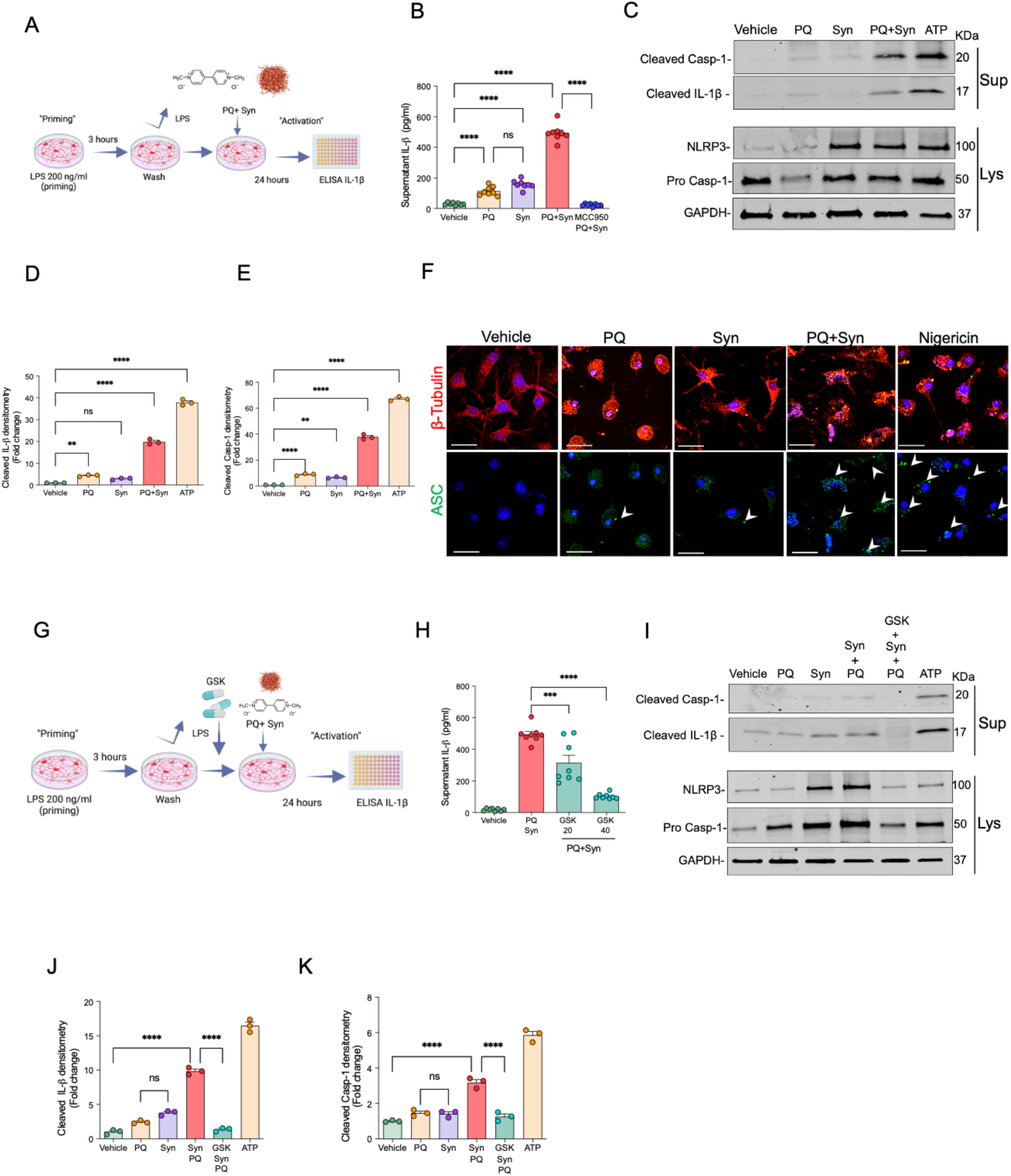
Paraquat enhances α-synuclein-mediated NLRP3 inflammasome activation in primary microglia. **(A)** Schematic of the experimental paradigm: LPS-primed primary microglia were treated with paraquat (PQ, 100 μM), pre-formed α-synuclein fibrils (Syn, 5 μM), or both for 24 hours. **(B)** IL-1β secretion in LPS-primed microglia following treatment with PQ, Syn, PQ+Syn or pretreated with MCC950 (100nM) before PQ+Syn. ATP (5 mM, 1 h) served as a positive control. **(C)** Immunoblot analysis of cleaved caspase-1 (p20) and cleaved IL-1β (p17) in supernatants (Sup), and NLRP3 and pro–caspase-1 in cell lysates (Lys) after treatment with PQ, Syn, or PQ+Syn, ATP (5 mM, 1 h) served as a positive control. **(D– E)** Densitometric quantification of cleaved caspase-1 (p20) and cleaved IL-1β (p17) from immunoblots. **(F)** Immunofluorescence staining of ASC (green) in LPS-primed microglia treated with PQ, Syn, or PQ+Syn, showing ASC speck formation (white arrows). Nigericin (10 μM) served as a positive control. Nuclei were counterstained with DAPI (blue). Scale bar: 20 μm. **(G)** Schematic of the experimental paradigm for NOX2 inhibition: LPS-primed microglia were pretreated for 30 min with the NOX2 inhibitor GSK2795039 prior to PQ+Syn exposure. **(H)** IL-1β secretion in primary microglia treated with PQ+Syn in the presence of GSK2795039 (20 or 40 μM). **(I)** Immunoblot analysis of cleaved caspase-1 (p20 and cleaved IL-1β (p17) in supernatants, and NLRP3 and pro–caspase-1 in lysates following PQ+Syn treatment with or without GSK2795039 (40 μM). Data are presented as mean ± SEM from ≥3 independent experiments. Statistical analysis was performed using one-way ANOVA with Tukey’s post hoc test; *p < 0.05, ***p < 0.001, ****p < 0.0001; ns, not significant.

Immunostaining for ASC revealed minimal speck formation with αSyn alone, while paraquat induced robust ASC specks. Paraquat+αSyn co-treatment led to widespread ASC speck formation and microglial swelling and rupture, consistent with pyroptosis (Fig. 2F). Nigericin, used as a positive control, also induced inflammasome mediated ASC speck formation. To confirm the role of NOX2 (Miller et al., 2007; Taetzsch & Block, 2013), we treated cells with GSK2795039 (20 and 40 μM) after LPS priming (Fig. 2G). GSK2795039 dose-dependently reduced IL-1β release in paraquat+αSyn-treated cells (Fig. 2H) and suppressed cleaved IL-1β and caspase-1 levels (Fig. 2I–K). Together, these findings indicate that paraquat amplifies αSyn-induced NLRP3 inflammasome activation via NOX2-derived ROS.

### Paraquat induces pyroptosis in a dose-dependent manner and synergizes with α-synuclein in primary microglia

Given that PQ+Syn-treated microglia displayed a swollen, ruptured morphology with ASC speck formation and release (Fig 2F), hallmarks of pyroptotic cell death (Fernandes-Alnemri et al., 2007), we investigated whether paraquat induces pyroptosis. A caspase-1-dependent LDH release assay revealed a dose-dependent increase in pyroptotic cell death following treatment with paraquat (Fig. 3A) similar to nigericin (10 μM, 1 h), with the caspase-1 inhibitor VX-765 (20 μM) suppressing cell lysis (Fig 3A).

**Figure 3.**
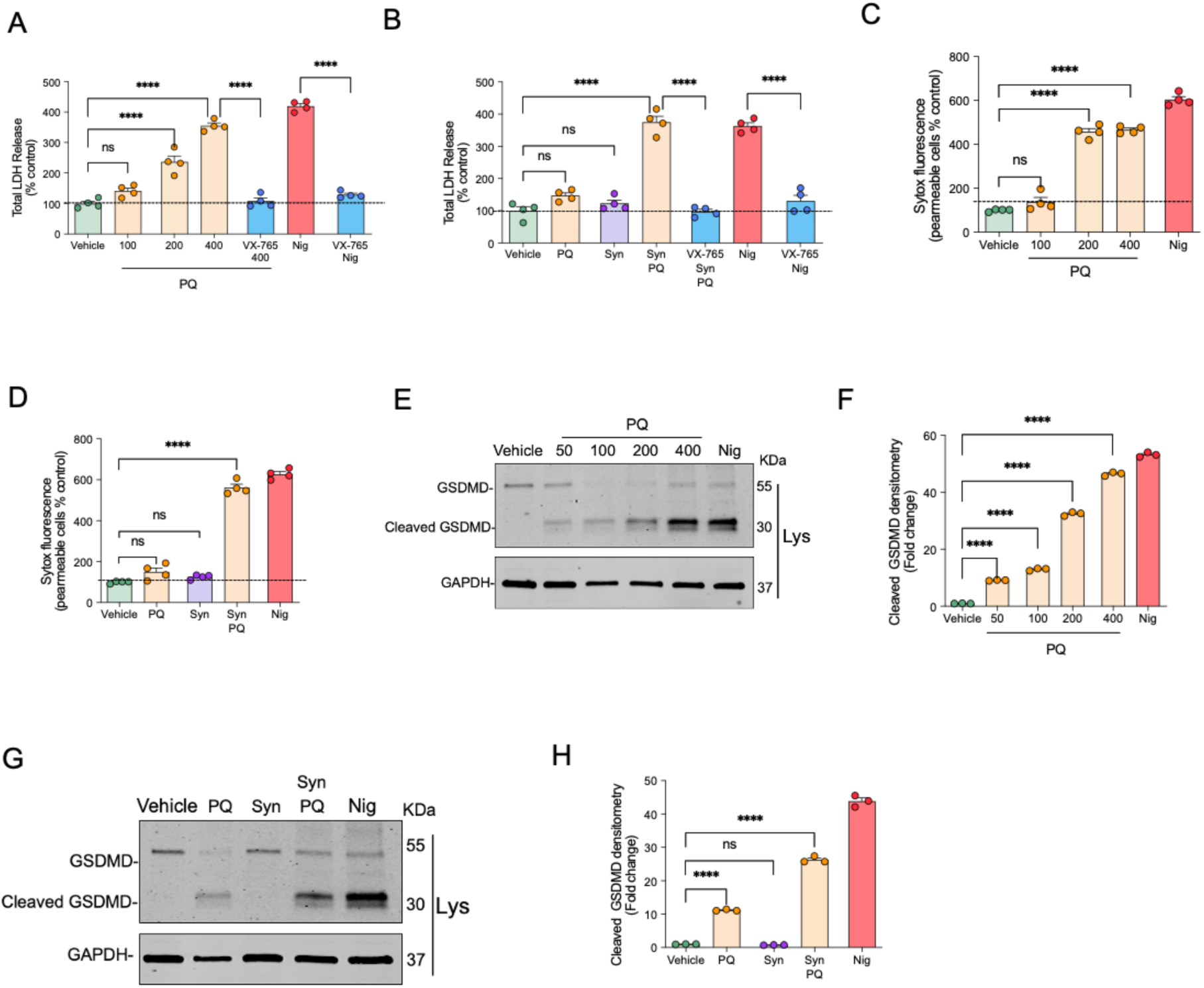
Paraquat induces pyroptosis in a dose-dependent manner and in synergizes with α-synuclein in primary microglia (A-B) Quantification of pyroptotic cell death by caspase-1 dependent LDH release assay in LPS-primed primary microglia treated (A) with increasing concentrations of paraquat (PQ, 100–400 μM) or (B) with PQ (100 μM), pre-formed α- synuclein fibrils (Syn, 5 μM), or both (PQ+Syn). Nigericin (10 μM) served as a positive control. Caspase-1 dependence was confirmed by the caspase-1 inhibitor VX-765 (20 μM) **(C- D)** SYTOX green uptake assay for membrane permeability following treatment with (C) PQ (100–400 μM) or (D) PQ, Syn, or PQ+Syn. Data are shown as fold change over vehicle-treated controls. **(E)** Immunoblot analysis of full-length and cleaved gasdermin D (GSDMD) in lysates from microglia treated with PQ (50–400 μM); nigericin (10 μM) was used as a positive control. **(F)** Densitometric quantification of cleaved GSDMD from immunoblots in (E). **(G)** Immunoblot analysis of GSDMD and cleaved GSDMD in lysates from microglia treated with PQ (100 μM), Syn (5 μM), or PQ+Syn. **(H)** Densitometric quantification of cleaved GSDMD from immunoblots in (G).Data represent mean ± SEM from ≥3 independent experiments. Statistical analysis was performed using one-way ANOVA with Tukey’s post hoc test; *p < 0.05, ****p < 0.0001; ns, not significant.

Neither paraquat (100 μM) nor αSyn (5 μM) alone significantly induced LDH release. However, their combination led to robust pyroptosis, comparable to the nigericin positive control, with cell lysis in each case abrogated by VX-765 (Fig. 3B). Similarly, SYTOX green uptake showed increased membrane permeability at paraquat doses ≥200 μM, with the PQ+αSyn combination significantly increasing permeability (Fig. 3C–D). Western blot analysis confirmed dose-dependent upregulation of cleaved gasdermin D (GSDMD), the key initiator of inflammasome-dependent pyroptosis (He et al., 2015) with paraquat treatment (Fig. 3E–F). PQ+αSyn co-treatment markedly increased cleaved GSDMD levels compared to αSyn alone, which showed no detectable cleavage (Fig. 3G–H), confirming paraquat-induced pyroptosis in the presence of αSyn.

### Paraquat activates the NLRP3 inflammasome and triggers pyroptosis in human microglia-like cells via a NOX-2 derived ROS

To confirm the clinical relevance of our findings, we investigated whether paraquat also activates the NLRP3 inflammasome in human microglia. Human monocyte-derived microglia-like cells (MDMi) were generated from healthy donor blood, and differentiated over 14 days using a defined cytokine cocktail, following established protocols (Albornoz, Amarilla, et al., 2023; Ryan et al., 2017) (Fig.4A). The resulting MDMi expressed key microglial markers TMEM119 and P2RY12 (Fig.4B).

**Figure 4.**
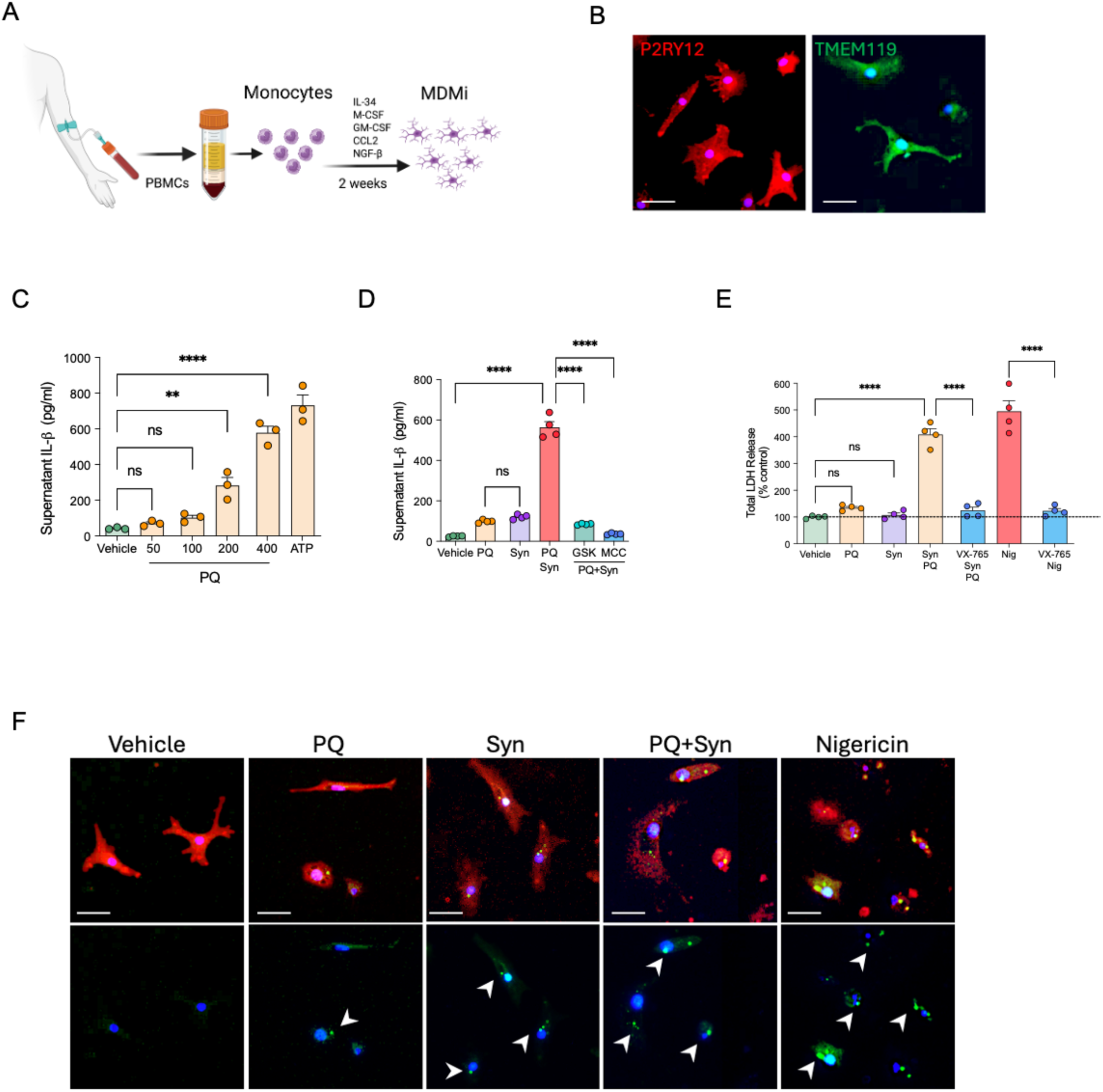
Paraquat activates the NLRP3 inflammasome and induces pyroptosis in human microglia via a NOX-2 derived ROS. **(A)** Schematic of the experimental workflow for generating human monocyte-derived microglia-like cells (MDMi) from peripheral blood mononuclear cells. **(B)** Immunofluorescence staining of microglial markers P2RY12 (red) and TMEM119 (green) confirming microglial identity. Nuclei were counterstained with DAPI (blue). Scale bar: 20 μm. **(C)** Dose-dependent IL-1β secretion in LPS-primed MDMi treated with increasing concentrations of paraquat (PQ, 50–400 μM) for 24 h. ATP (5 mM, 1 h) was used as a positive control for inflammasome activation. **(D)** IL-1β secretion in MDMi treated with PQ (100 μM), α-synuclein fibrils (Syn, 5 μM), or both (PQ+Syn). Cells were pretreated with the NOX2 inhibitor GSK2795039 (GSK, 40 μM) or the NLRP3 inhibitor MCC950 (100 nM).**(E)** LDH release assay to quantify pyroptotic cell death in MDMi treated with PQ, Syn, or PQ+Syn. Nigericin (10 μM) was used as a positive control. Caspase-1 dependence was confirmed by pretreatment with the caspase-1 inhibitor VX-765 (20 μM).**(F)** Immunofluorescence staining for ASC (green) in LPS-primed MDMi treated with PQ, Syn, or PQ+Syn, showing characteristic ASC speck formation (white arrows). Nuclei were counterstained with DAPI (blue). Nigericin was used as a positive control. Scale bar: 20 μm. Data represent mean ± SEM from ≥3 independent donors. Statistical analysis was performed using one-way ANOVA with Tukey’s post hoc test; *p < 0.05, ****p < 0.0001; ns, not significant.

Paraquat induced a dose-dependent increase in IL-1β release (Fig. 4C). PQ+αSyn co-treatment significantly enhanced IL-1β levels compared to individual treatments, and this effect was blocked by GSK2795039 and completely abolished by MCC950 (Fig. 4D), indicating that a NOX2–ROS–NLRP3 signalling axis underpins IL-1β release. Caspase-1–dependent LDH release assays showed that neither PQ nor αSyn alone induced significant pyroptosis, but their combination triggered robust LDH release that was abrogated by VX-765 (Fig. 4E). Immunostaining confirmed ASC speck formation and microglial rupture with paraquat+αSyn, consistent with pyroptotic cell death (Fig. 4F). Together, these findings demonstrate for the first time that PQ triggers NOX2-derived ROS to activate the NLRP3 inflammasome and ensuing pyroptosis in human microglia-like cells exposed to fibrillar αSyn.

## Discussion

Our study provides novel insights into the interplay between paraquat, αSyn, and the microglial NLRP3 inflammasome, offering a potential mechanistic explanation for how environmental toxins may exacerbate αSyn-driven neuroinflammation in PD. While paraquat has long been associated with increased PD risk through oxidative stress and dopaminergic toxicity (Costello et al., 2009; Sharma & Mittal, 2024; Tanner et al., 2011), our data now suggest that this compound also primes a highly inflammatory form of programmed death—pyroptosis—in microglia, particularly in the presence of αSyn aggregates.

Critically, we show that paraquat triggers NLRP3 inflammasome activation in LPS-primed microglia via a ROS-dependent pathway. This effect was abolished in NLRP3 knockout cells, implicating the canonical NLRP3-ASC-caspase-1-GSDMD axis in the observed pyroptotic response. These findings are consistent with earlier reports of paraquat-mediated inflammasome activation in peripheral macrophages (Z. Liu et al., 2015), but our data expand this mechanism to CNS-resident microglia under neurodegenerative conditions. This is particularly important given recent re-evaluations of paraquat’s CNS penetration and persistence, which suggest the compound can cross the blood-brain barrier under inflammatory conditions or with chronic exposure (Breckenridge et al., 2013).

In line with our previous work (Albornoz, Gordon, et al., 2023; Gordon et al., 2018), we confirm that αSyn can activate sublytic NLRP3 signalling in microglia, resulting in IL-1β release. However, the combination of paraquat and αSyn induced a robust pyroptotic phenotype, evidenced by ASC speck formation, caspase-1 activation, GSDMD cleavage, and caspase-1-dependent LDH release. This highlights a previously underappreciated synergistic interaction, wherein paraquat-induced ROS may act as a secondary signal that facilitates αSyn-driven inflammasome assembly and pyroptosis execution.

This cooperative mechanism aligns with recent data emphasizing αSyn–microglia crosstalk in disease initiation. αSyn fibrils activate NLRP3 microglia *in vivo*, accelerating dopaminergic neurodegeneration in a manner dependent on inflammasome components (Huang et al., 2023). Moreover, αSyn-induced microglial reactivity is sufficient to initiate neurodegenerative cascades even in the absence of overt neuronal αSyn pathology (Sirerol-Piquer et al., 2025). Our findings reinforce this paradigm; environmental triggers such as paraquat can exacerbate αSyn-mediated microglial inflammasome responses, thereby lowering the threshold for neuroinflammation and pyroptosis.

The translational relevance of our study extends beyond the bench. Paraquat remains in use in numerous countries, including Australia, despite mounting evidence of its neurotoxicity. Our findings carry significant public health implications for communities with occupational or incidental paraquat exposure, particularly in agricultural settings. This is underscored by a recent population-based cohort study (Krzyzanowski et al., 2025), which found that individuals living near golf courses, where herbicides including paraquat are commonly applied, had a significantly elevated risk of developing PD. This strengthens the call for re-evaluating environmental risk factors in PD and reinforces the urgency of regulatory decisions concerning paraquat in countries where it is still permitted.

Our data also emphasize the need for vigilance in countries like Australia, where agricultural workers and nearby residents may face chronic low-level exposure to paraquat. Given the compound’s persistence and the long prodromal phase of PD, understanding how such exposures interact with individual susceptibility (e.g., αSyn pathology, inflammasome priming) is critical for preventive strategies.

While our study provides mechanistic evidence of paraquat–αSyn synergy in microglial pyroptosis, several limitations warrant consideration. Chief among these is the *in vitro* nature of our model. Although both murine and human microglia exhibited inflammasome activation, the cellular microenvironment *in vitro* cannot fully replicate the complexity of *in vivo* neuroinflammation, nor the dynamic interactions among neurons, glia, and peripheral immune cells. Furthermore, our reliance on LPS priming, although a standard inflammasome model, may not entirely capture the physiological triggers present in PD brains.

To address these gaps, we propose future *in vivo* investigations incorporating dual exposure paradigms of paraquat and αSyn (e.g., fibrils or overexpression models), ideally in *Nlrp3*-competent versus *Nlrp3-*deficient backgrounds. These studies would establish whether the synergistic inflammasome-mediated neurotoxicity observed here translates to progressive dopaminergic neurodegeneration and motor impairment *in vivo*. This would also clarify whether paraquat accelerates or intensifies αSyn pathology spread and whether microglial pyroptosis contributes causally to neuronal dysfunction.

This study contributes to the growing body of literature on pyroptosis in the brain. The role of GSDMD-mediated cell death in neuroinflammation is increasingly recognized, with studies demonstrating its presence in both human and murine models of neurodegeneration (Hu, Wang, Li, & Yang, 2022; X. Liu, Xia, Zhang, Wu, & Lieberman, 2021). Pyroptosis may exacerbate neurotoxicity not only by releasing proinflammatory cytokines but also through cellular lysis and release of damage-associated molecular patterns (DAMPs), thus perpetuating a feedforward inflammatory loop.

In conclusion, we propose that paraquat could potentially act as a mechanistic environmental co-factor in PD by promoting microglial NLRP3 inflammasome activation and pyroptotic cell death, particularly in the context of αSyn pathology. These findings underscore the urgent need for inflammasome-targeted therapeutics in neurodegenerative diseases driven by environmental exposures.

## Author Contributions

EA conceived the project and designed experiments with TW and MC, KS, and AR assisting in provision of NLRP3 inhibitor reagents and *Nlrp3* knockout mice. YC, VD, NK and EA performed the experiments and data analysis. EA and YC wrote the manuscript. All authors edited and approved the final version of the manuscript.

